# Superparamagnetic iron oxide enclosed hollow gold nanostructure with tunable surface plasmon resonances to promote near-infrared photothermal conversion

**DOI:** 10.1101/2022.02.07.479434

**Authors:** Muzhaozi Yuan, Xuhui Feng, Tian-Hao Yan, Jingfan Chen, Xuezhi Ma, Preston Cunha, Shoufeng Lan, Ying Li, Hong-Cai Zhou, Ya Wang

**Affiliations:** J. Mike Walker ‘66 Department of Mechanical Engineering, Texas A&M University, College Station, TX 77840 United States; Department of Chemistry, Texas A&M University, College Station, TX 77843 United States; Department of Materials Science & Engineering, Texas A&M University, College Station, TX 77840 United States; Department of Biomedical Engineering, Texas A&M University, College Station, TX 77843 United States; Department of Electrical and Computer Engineering, Texas A&M University, College Station, TX 77843 United States

**Keywords:** SPIO-HGNS, near-infrared, tunable wavelength, photothermal, pulsed laser

## Abstract

In this study, to enhance deep tissue penetration by near-infrared (NIR) light, a novel superparamagnetic iron oxide enclosed hollow gold nanoshell (SPIO-HGNS) structure with tunable size and surface plasmon resonance (SPR) in the NIR range was designed and synthesized through a 2-step template-enabled galvanic replacement reaction. Here, Ag coated SPIO (SPIO-Ag) was prepared as a template with tunable outer diameters by way of adjusting the Ag content. SPIO-HGNS with variable hollow gold inner diameters can then be synthesized based on the determined outer diameter of the SPIO-Ag template through a galvanic replacement reaction between HAuCl_3_ and Ag coating on the SPIO surface. With incrementing amounts of Ag, three SPIO-HGNS structures were synthesized with comparable shell thicknesses around 6.7 nm and an average inner diameter of 38.7, 39.4, and 40.7 nm, respectively, evidenced by TEM and ICP results. The structure of SPIO-HGNS was confirmed by identifying Au111 lattice and the elemental mapping of Fe and Au using energy-dispersive X-ray spectroscopy. The Ultraviolet-Visible-NIR absorption spectra showed red-shifted SPR peaks (820, 855, and 945 nm) with the increasing inner diameters of SPIO-HGNS, which was also supported by an absorption cross-section simulation. The photothermal results showed that the three SPIO-HGNS structures, when exposed to ~30 s of 400 mW laser irradiation, exhibited photothermal temperature rises of 5.9, 4.6, and 2.9 °C, respectively. This study explored the tuning of SPR properties in NIR-responsive magneto-plasmonic nanoparticles through a facile preparation procedure, paving the way for potential applications in photothermal therapies.

## 1. Introduction

The unique optical properties of plasmonic nanoparticles (NPs) make them attractive in many biomedical applications. It is reported that plasmonically generated heat can activate temperature-sensitive channels in neurons [1] for the excitation [2] or inhibition [3] of action potentials. Additionally, it can regulate Ca^2+^ levels [4], which is critical for neuron communication [5, 6] and the regulation of cytoskeleton growth behaviors via phosphorylation [7]. Our previous work demonstrated that green light stimulated superparamagnetic iron oxide core-gold shell (SPIO-Au) NPs promote neuronal differentiation and neurite outgrowth via plasmonic heating [8]. However, the SPR peak of the plasmonic NPs is located at the visible light region, which has very low tissue penetration ability and thus limits the NPs biomedical applications. To address these limitations, the coupling of upconversion NPs with plasmonic NPs has been explored as a means to upconvert the incident near-infrared (NIR) light to visible light. This is because NIR light has significantly stronger tissue penetration ability than visible light [9]. The photothermal effects can be induced by the visible light emitted from the upconversion NPs, which overlaps with the plasmonic resonance frequency of plasmonic NPs [10, 11, 12]. Several materials have been demonstrated to be plasmonically stimulated by upconversion NPs—including Au nanorods, Au nanoshells, Au clusters, and CuS [11]. However, due to the very low light absorption coefficient of upconversion NPs, the design of upconversion-plasmonic NPs with high quantum yields remains challenging, limiting the photothermal performance of plasmonic NPs [13]. Additionally, the synthesis procedures of upconversion-plasmonic NPs are limited by their complexity. To develop simplified synthesis procedures and enhance photothermal performance, it is necessary to develop new structurally modified plasmonic NPs with tunable and strong SPR peaks in the NIR region.

Recently, hollow gold nanoshell (HGNS) has demonstrated tunable SPR response from the visible light to NIR regions [14]. This HGNS structure is composed of a spherical thin gold shell filled with an aqueous medium [15]. Previously synthesized HGNS structures have SPR peaks ranging from 735 nm to 828 nm [16, 17, 18, 19]. Zhang *et al*. were able to tune the SPR peak of HGNS from 545 nm to 740 nm by controlling the inner and outer diameters of HGNS [20]. Xie *et al*. synthesized HGNS with an SPR peak longer than 1000 nm [21]. These results indicate that the plasmonic properties of HGNS highly depend on its size and thickness. Although these HGNS structures are NIR stimulable, they lack the targeting abilities that would enable selective delivery to the area of interest for photothermal tumor ablation. Compared to other targeting strategies that rely on local microenvironment stimuli and may cause off-target localization, magnetic targeting can be more selective and efficient by the accumulation of NPs through external magnetic field guidance with minimal side-effects [22, 23]. Therefore, to enable the selective targeting ability of HGNS, magnetic NPs can be combined with HGNS. Among the available magnetic materials, superparamagnetic iron oxide (SPIO) NPs have been identified as the most promising option due to their high magnetization properties and small size (10-15 nm) [24]. In fact, SPIO NPs have been used extensively in biomedical fields such as bioimaging [25, 26, 27] and targeted drug/gene delivery [28, 29].

Several attempts have been made to prepare SPIO enclosed HGNS (SPIO-HGNS) nanostructures. Traditional methods employed a direct seed-growth coating method to form a gold shell at the surface of SPIO using hydroxylamine as a reductant [30]. However, due to the size limit of SPIO cores (10-15 nm), the plasmonic peak of the complex is restricted below 650 nm, and cannot be tuned to the NIR window [30, 31, 32]. To address this limitation, a large size of HGNS can be designed and prepared by forming an additional layer on the SPIO surface to enlarge the size of the template before the growth of the gold layer. For example, Ji *et al*. [33] and Melancon *et al*. [34, 35] applied tetraethyl orthosilicate (TEOS) and coated a SiO_2_ layer on SPIO, with the HGNS formation later being initiated on the SiO_2_ surface to form an SPIO/SiO_2_/HGNS nanostructure. The prepared nanostructure exhibited a wide light absorption ranging from 500 nm to 1100 nm and was centered at ~800 nm. Han *et al*. [36] later reported an updated preparation method by applying polypyrrole as the binder to enhance the surface affinity between the TEOS and SPIO. However, these methods add multiple steps to their experimental procedures that increase complexity and time-cost to form SPIO-HGNS nanostructures. Further, these methods are unable to realize the tuning of the SPR peak of SPIO-HGNS and maximization of the absorption of SPIO-HGNS at specific NIR wavelengths.

In this work, to solve the aforementioned challenges, we proposed a one-pot preparation method for SPIO-HGNS structures with tunable light absorption properties. We hypothesized that, during the galvanic replacement reaction, the HGNS size can be controlled by varying the size of core-shell Ag coated SPIO (SPIO-Ag) NP templates. First, with the existence of a Ag^+^ precursor solution, NH_2_OH∙HCl was used to promote the formation and growth of an Ag layer at the surface of SPIO seeds, instead of Ag nucleation in bulk solution. During this step, the outer diameter of the SPIO-Ag intermediate nanostructures was controlled by varying the concentration of the Ag^+^ precursor. Then the SPIO-Ag NPs were applied as templates for the formation of HGNS through galvanic replacement reaction. During this reaction, the Ag layer was eroded by HAuCl_4_ and HGNS was formed [37]. SPIO NPs were enclosed inside the HGNS to form an SPIO-HGNS structure, as shown in Fig. 1. Transmission electron microscopy (TEM) was used to reveal the size and structure of SPIO-HGNS and confirm SPIO-HGNS formation. The light absorption spectra of SPIO-HGNS NPs were measured to identify the relationship between the size of NPs and the position of the SPR peak. Photothermal conversion tests under ultrafast laser excitation, in conjunction with a finite element model (FEM) simulation, were conducted to determine the photothermal conversion efficiency of SPIO-HGNS at different sizes. Additionally, a boundary element method was applied to further numerically identify the relationship between the SPR peak position and the size of SPIO-HGNS NPs.

**Fig. 1.**
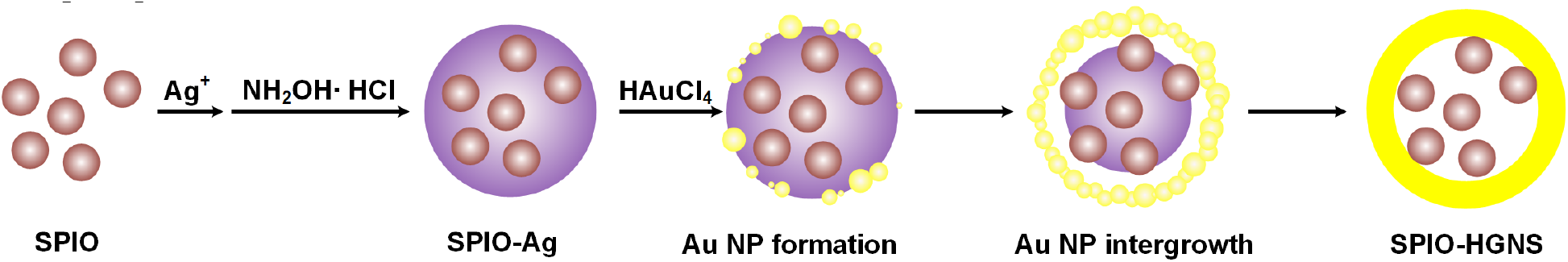
Schematic illustration showing the preparation process of SPIO-HGNS NPs.

## 2. Experimental

### 2.1 Chemicals and materials

SPIO NPs (EMG 304, Ferrotec, Santa Clara, CA), trisodium citrate (Na_3_C_6_H_5_O_7_, 99%, Alfa Aesar, Haverhill, MA), silver nitrate (AgNO_3_, ≥99.0%, Sigma-Aldrich, St. Louis, MO), sodium hydroxide (NaOH, ≥98.0%, Sigma-Aldrich, St. Louis, MO), hydroxylamine hydrochloride (NH_2_OH∙HCl, 99+%, Acros Organics, Fairlawn, NJ), and chloroauric acid (HAuCl_4_, ≥99.0%, Sigma-Aldrich, St. Louis, MO).

### 2.2 Preparation of SPIO-HGNS NPs

To prepare the SPIO-HGNS structures, 1 ml of 0.64 mg/L SPIO NPs in deionized (DI) water, 1 ml of 30 mmol/L trisodium citrate (Na_3_C_6_H_5_O_7_, Alfa Aesar, 99%) solution, and a specified amount of DI water was mixed in a rounded bottom flask, which was placed in an oil bath maintained at 60 °C. The solution was stirred for 10 min, which was followed by the addition of the desired amount of 20 mmol/L AgNO_3_ (Sigma-Aldrich, ≥99.0%) solution (1.0 ml, 1.5 ml, and 2.0 ml for SPIO-HGNS-I, II, and III, respectively) into the liquid mixture. Then the solution was stirred for another 5 min. A 5 mmol/L NaOH (Sigma-Aldrich, ≥98.0) solution was then gradually added into the mixture to adjust the pH value of the solution to ~10. After 30 min of stirring, 0.25 ml of 200 mmol/L NH_2_OH∙HCl (Acros Organics, 99+%) solution was added into the round-bottom flask. After another 3.5 h of stirring, 5 mmol/L NaOH solution was added into the reaction solution to adjust the pH to ~6.0, a pre-calculated volume (0.4×volume of AgNO_3_ solution) of 25 mmol/L HAuCl_4_ (Sigma-Aldrich, ≥99.0%) solution was then added into the reaction solution which was stirred for another 30 min. After the reaction, the SPIO-HGNS NPs were separated by centrifugation and rinsed with 1.2 mmol trisodium citrate solution 4 times to remove the excess uncoated SPIO. The NPs were then redispersed into a 1.2 mmol trisodium citrate solution and stored at 4 °C.

### 2.3 Material characterization

The structure and morphology of the SPIO-HGNS NPs were analyzed on an FEI Tecnai G2 F20 ST field emission transmission electron microscope (FE-TEM), operated at an accelerating voltage of 200 kV. The high-resolution scanning electron microscopy (SEM) image of the SPIO-HGNS NPs was taken on a JEOL JSM-7500F ultra high-resolution field emission scanning electron microscope equipped with a high brightness conical FE gun and a low aberration conical objective lens. The size distribution of the SPIO-HGNS NPs was analyzed by measuring a minimum of 50 NPs using ImageJ. The inner and outer diameters for each NP were measured using ImageJ and the difference between them was calculated as shell thickness. The inner diameter and shell thickness of SPIO-HGNS were expressed as average values and standard deviations. The light absorption spectra of SPIO-HGNS were measured using a Hitachi U-4100 UV-Vis-NIR spectrophotometer (500-1200 nm). In a typical measurement, 3 ml of NP solution was added to a quartz cuvette.

To determine the elemental composition of SPIO-HGNS, inductively coupled plasma mass spectroscopy (ICP-MS) analysis was conducted on a PerkinElmer NexION 300D ICP mass spectrometer using 2% nitric acid and 1% hydrochloric acid as the analytical matrix. In detail, the SPIO-HGNS NPs were digested by aqua regia at 80 °C overnight. After cooling to room temperature, the sample solutions were diluted with 2% nitric acid and 1% hydrochloric acid. The triplicate results were averaged to determine the metal concentrations.

### 2.4 Photothermal conversion efficiency evaluation

To evaluate the photothermal conversion efficiency, a previously developed experimental set-up was adopted to measure the temperature profile of SPIO-HGNS NPs dispersed in DI water and phosphate-buffered saline (PBS) respectively [38, 39]. A Coherent Chameleon series ultrafast laser generator (operation power: 400 mW, pulse frequency: 80MHz, pulse duration: 140 fs) was used as the light source. A polymethyl methacrylate semi-micro cuvette was used as the container, and 0.8 ml of sample solution was used in each test. A thermocouple (Omega, Stamford, CT, K-type, 0.076 mm wire diameter, and 0.33 mm bead diameter) was used to measure the temperature variation vs. time under laser beam irradiation. A National Instruments NI9219 control system was employed to record the temperature vs. time curve, which was incorporated into a previously developed FEM model to simulate the temperature rise of each SPIO-HGNS solution [39]. Photothermal conversion efficiencies were obtained during this process by matching the modeled temperature rise with the experimental data.

## 3. Results and discussion

### 3.1 Material structure characterization

In our previous work [32, 38, 40], we synthesized the SPIO-Au core-shell NPs with an SPR peak at the visible light region. In this work, to further modify the structure and tune the SPR peak to the NIR region, we first synthesized Ag coated SPIO NPs. The existence of hydroxylamine hydrochloride allows the formation of Ag coating on SPIO NPs and the thickening of the coated Ag layer instead of the formation of free Ag NPs in the bulk solution. Then the addition of HAuCl_4_ initiates the galvanic replacement reaction, where Ag is eroded by HAuCl_4_ and HGNS is formed. This leads to the nanostructure of SPIO-enclosed HGNS, denoted as SPIO-HGNS in this study.

Fig. 2 shows the TEM images of the synthesized SPIO-HGNS-I, II, and III NPs. A NP size survey, based on several TEM images that include > 50 NPs for each of the 3 samples, was conducted. The inner diameter and shell thickness of SPIO-HGNS were calculated and listed in Table 1. The trend that inner diameter increases with a higher Ag ratio matches the proposed synthetic mechanism that a higher level of Ag^+^ amount leads to the formation of a larger Ag layer and a larger inner diameter of SPIO-HGNS. The parallel lattice fringes of 0.24 nm spacing (Fig. 2(b)), along with the inset of fast Fourier transformations (FFT), confirms the presence of Au111 lattice. However, the lattice of SPIO cannot be observed, which may be due to blocking by the high-contrast Au element as SPIO is wrapped by HGNS. The high-resolution SEM image, as shown in Fig. S3, confirmed the uniform morphology of SPIO-HGNS NPs.

**Table 1.** The measured inner diameter and shell thickness for SPIO-HGNS samples.

|  | SPIO-HGNS-I | SPIO-HGNS-II | SPIO-HGNS-III |
| --- | --- | --- | --- |
| Measured HGNS shell inner diameter from TEM (nm) | $38.7 \pm 9.9$ | $39.4 \pm 11.8$ | $40.7 \pm 11.1$ |
| Measured HGNS shell thickness from TEM (nm) | $6.7 \pm 1.3$ | $6.5 \pm 1.1$ | $7.0 \pm 2.1$ |

**Fig. 2.**
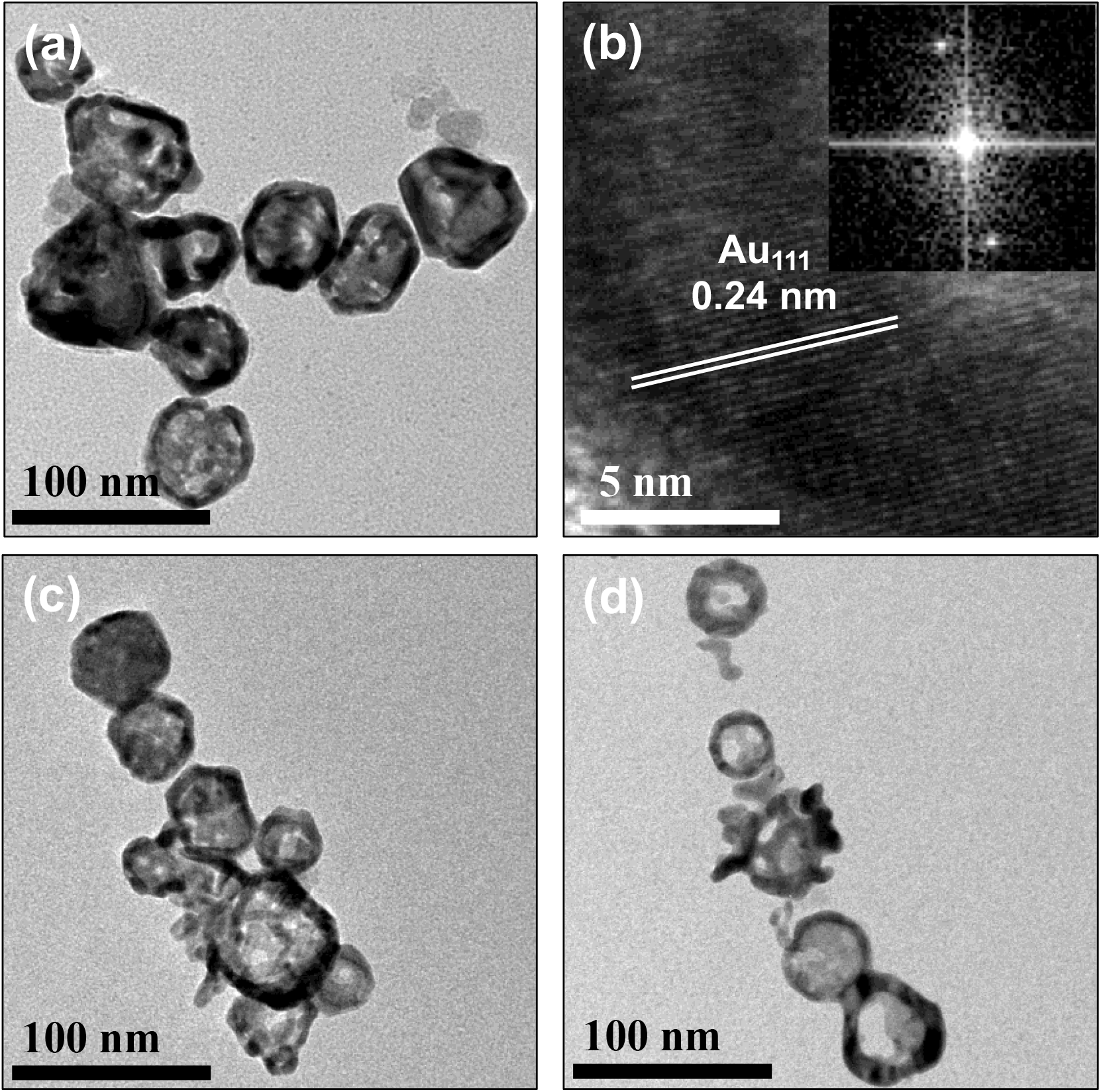
TEM images of SPIO-HGNS NPs. (a) SPIO-HGNS-I; (b) Lattices and FFT of SPIO-HGNS-I; (c) SPIO-HGNS-II; (d) SPIO-HGNS-III. More TEM images of SPIO NPs and SPIO-HGNS NPs, including concentrated samples, are listed in Fig. S1. The size distribution of each type of NP based on >50 NPs is listed in Fig. S2.

To further confirm the hollow structure of the as-synthesized NPs, a TEM image and a scanning transmission electron microscope (STEM) image using high-angle annular dark-field (HAADF) of a single SPIO-HGNS-I NP were taken, as shown in Fig. 3(a) and (b). The core-shell structure of the SPIO-HGNS-I NPs was verified by the elemental mapping with the energy-dispersive X-ray spectroscopy (EDS). As shown in the elemental mapping images in Fig. 3(c) and (d), gold (Au) is distributed throughout the whole NP. Iron (Fe), even with the background noise from the TEM specimen grid, is more concentrated in the center of the NP with lower signal intensity. This proves that SPIO (contains Fe) NPs are wrapped by HGNS. The EDS results also indicate that the Fe element is widely distributed in the hollow space, a sign that there are multiple SPIO NPs wrapped by HGNS, as shown in Fig. 3d. To further narrow down the size distribution and improve the morphology of SPIO-HGNS, our future work will focus on parametric study of pH value, the reactants’ concentration and the addition of surfactants, etc.

**Fig. 3.**
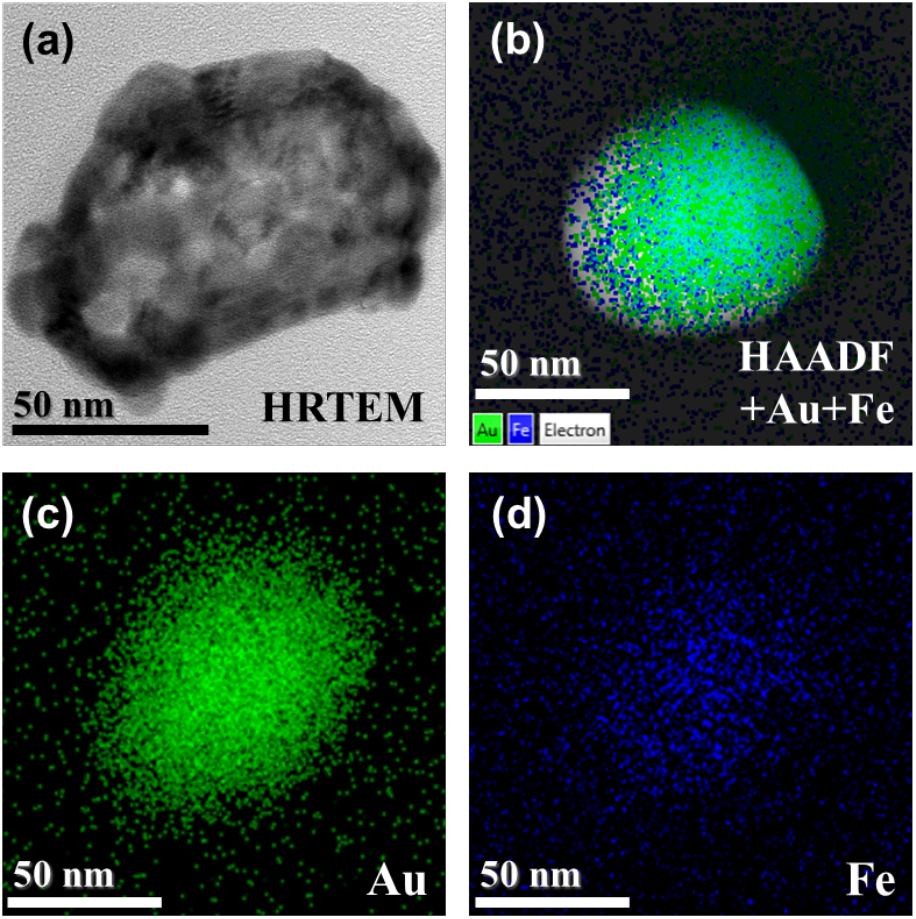
(a) HRTEM image of SPIO-HGNS-I; (b) HAADF image and TEM-EDS analysis of SPIO-HGNS-I; (c)-(d) TEM-EDS data of Au and Fe, suggesting the formation SPIO enclosed hollow gold nanostructure.

The elemental analysis was performed using ICP-MS. As shown from Table S1, the existence of both Fe and Au elements in the SPIO-HGNS-I, II, and III samples are confirmed, further demonstrating the coexistence of both SPIO core and gold shells. The mole ratio of Au/Fe is estimated to be 1.1, 2.2, and 5.9 for the SPIO-HGNS-I, II, and III samples, respectively. The increasing mole ratio is in accordance with the increasing inner diameter of the HGNS shell as well as the increasing Au precursor amount added to the reaction solution during the sample preparation process.

The modification of SPIO-HGNS by polymers was performed to examine the stability of SPIO-HGNS NPs. By functionalizing SPIO-HGNS with positively charged thiol ligands, the zeta potential was changed from −44.5 mV to 30.3 mV, and the hydrodynamic diameter was changed from 211.8 nm to 160.8 nm (Table S2). This suggests the effect of polymer coating in increasing the dispersity and enhancing the stability of SPIO-HGNS NPs. The positive zeta potential of Au NPs is usually favored for efficient intracellular delivery without toxic effects at low concentrations. This is due to higher cellular uptake efficiency through direct diffusion and stronger interactions between freely dispersed NPs and cell organelles [40, 41, 42]. Also, in our previous work, PEG-coated SPIO-Au NPs have shown excellent biocompatibility in several cell lines for up to 80 ppm [38, 40]. In the future, we will further explore the biocompatibility of PEG-coated SPIO-HGNS NPs.

### 3.2 UV-Vis-NIR absorption and magnetic properties

To compare the plasmonic properties of SPIO-HGNS at different sizes, the UV-Vis-NIR absorbance spectra were measured. As shown in Fig. 4, SPIO-HGNS-I exhibits a wide light absorption in the range of 800-1200 nm with an SPR peak at ~ 820 nm, which is located in the NIR range. For SPIO-HGNS-II and SPIO-HGNS-III, with a higher Ag^+^ addition amount (20, 30, and 40 μmol Ag^+^ for SPIO-HGNS-I, II, and III, respectively) in the preparation process, the prepared nanostructures exhibit a red-shifted SPR peak. That is, SPIO-HGNS-II exhibits an SPR peak at ~855 nm, and SPIO-HGNS-III exhibits an SPR peak at ~945 nm. Since the peak is very broad, to further identify the overall redshift of the peak, we calculated the half-power bandwidth (1/√2 of the maximum peak position) of each spectrum. As shown in the inset of Fig. 4, the SPIO-HGNS-I, II, and III samples have a red-shifted half-power band of 627-1022 nm, 647-1081 nm and 736-1208 nm, respectively. The NIR light absorption properties of the SPIO-HGNS nanostructures are dictated by the combination of inner and outer diameters. As proved by Liang *et al*. [43] and Schwartzberg *et al*. [44], a larger shell inner diameter and/or a smaller outer diameter of HGNS led to a redshift in NIR absorption peak position. It was shown in Fig. 2 and Table 1 that the inner and outer diameters of HGNS were increased by using greater Ag^+^ amounts in the preparation procedure. This may likely lead to larger outer diameters of the SPIO-Ag templates and result in a change in both the inner and outer diameters of HGNS.

**Fig. 4.**
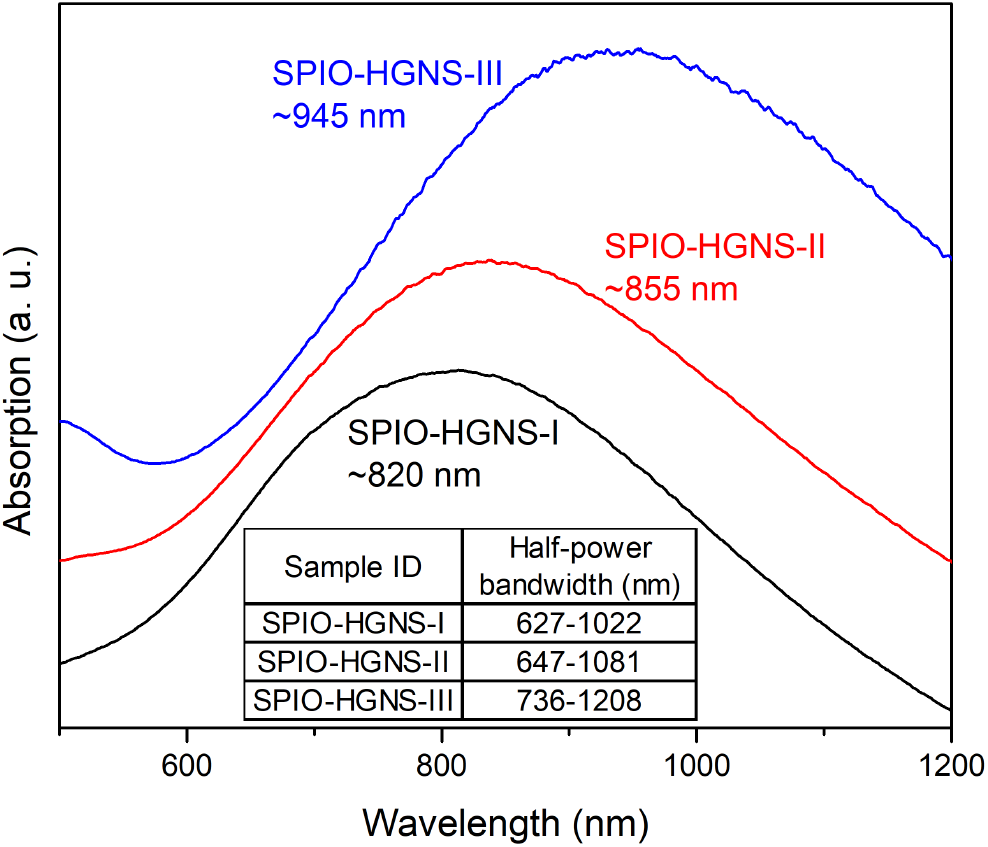
UV-vis-NIR absorption spectra of SPIO-HGNS at different size.

To confirm the magnetic properties of the as-prepared SPIO-HGNS NPs, the behavior of SPIO-HGNS-I under an external magnetic field was recorded. To simplify the test, SPIO-HGNS-I NPs were pre-concentrated by centrifugation at the bottom of a centrifugate tube. A piece of permanent magnet block was used to provide the external magnetic field. As shown in Fig. S4 (a), at 0 s, when the magnet is not applied, the SPIO-HGNS-I NPs stay at the bottom of the tube. It only takes ~5 s for the external magnetic field to attract almost all the SPIO-HGNS NPs towards the magnet block, as shown in Fig. S4 (b), which confirms the magnetic properties of the as-prepared SPIO-HGNS NPs.

### 3.3 Photothermal conversion performance

To evaluate the photothermal conversion capacity of SPIO-HGNS-I, the temperature profile measurements were performed on DI water as a baseline and SPIO-HGNS-I. The incident laser wavelength was set to be 820 nm to match the SPR peak of SPIO-HGNS-I. As shown in Fig. 5(a), in DI water (no SPIO-HGNS-I NPs), the laser merely causes an equilibrium temperature increase of ~2.5 °C after 400 s. By contrast, the SPIO-HGNS-I solution shows significantly higher temperature changes under laser irradiation. With an Au concentration of 38 ppm and 19 ppm, the SPIO-HGNS-I shows remarkable temperature rises of 29.5 °C and 26.0 °C, respectively, when reaching temperature equilibrium after 1000 s. Even with an Au concentration as low as 9 ppm, a considerable equilibrium temperature increase of 20.7 °C is found with SPIO-HGNS-I.

**Fig. 5.**
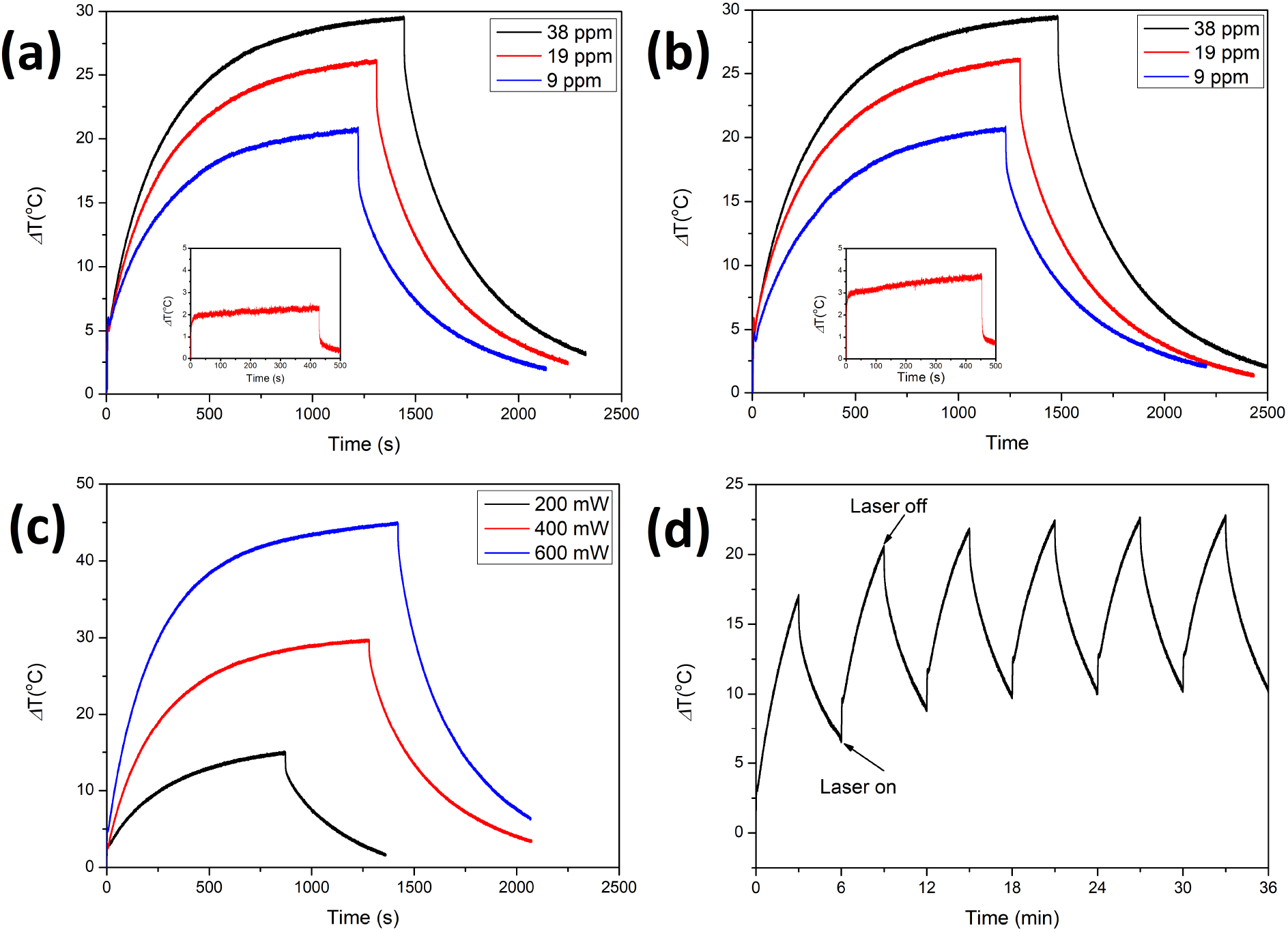
Temperature profile of SPIO-HGNS-I obtained (a) in DI water, (b) in PBS solution (c) in DI water under varied laser power from 200 mW to 600 mW, and (d) after six ON/OFF cycles of 820 nm laser irradiation at 400 mW. (ON: 3 min; OFF: 3 min) The inset of each figure shows the temperature profile of (a) DI water without NPs and (b) PBS solution without NPs.

To examine the biostability and photothermal conversion behavior of SPIO-HGNS-I in biological environments, a similar photothermal test was conducted on SPIO-HGNS-I dispersed in PBS, which was used to mimic the human-plasma environment. SPIO-HGNS-I was found to be stable in PBS without aggregation, suggesting high biostability in a bio-mimicking environment. As shown in Fig. 5(b), the photothermal conversion performance of SPIO-HGNS-I in PBS is similar to the results obtained in DI water (Fig. 5(a)) under the same laser irradiation setup: the maximum temperature increases of SPIO-HGNS-I at concentrations of 38 ppm, 19 ppm, and 9 ppm are 29.4 °C, 26.1 °C, and 20.6 °C, respectively. The blank test with pure PBS shows a negligible temperature rise compared with those obtained with the SPIO-HGNS-I solutions, indicating the superior photothermal conversion performance of SPIO-HGNS-I in a bio-mimicking environment.

To examine the change in temperature of SPIO-HGNS-I under varied levels of laser power, the temperature profile measurements were performed on SPIO-HGNS-I under a laser power of 200, 400 and 600 mW. As shown in Fig. 5(c), the higher maximum temperature rise was obtained with higher laser power. We observed that the temperature profiles of SPIO-HGNS NPs under different laser power intensities have no impact on the photothermal conversion efficiency, which is consistent with recent literature findings [45, 46].

Additionally, to evaluate the photothermal stability of SPIO-HGNS-I, six laser ON/OFF cycles were performed and the temperature profiles were recorded, as shown in Fig. 5(d). In each cycle, SPIO-HGNS-I NPs were irradiated by a 820 nm laser for 3 mins and then cooled down for 3 mins by turning off the laser. No significant reductions in the temperature rise were observed during these six cycles, suggesting outstanding photothermal stability of SPIO-HGNS-I NPs under the irradiation of the 820 nm laser.

To further investigate the relationship between the photothermal conversion of SPIO-HGNS NPs with varying size, the photothermal conversion tests were conducted on SPIO-HGNS-II and III dispersed in DI water. The laser wavelengths used for SPIO-HGNS-II and III were 855 nm and 945 nm, respectively, to match the maximum SPR peak of each sample. As shown in Fig. 6, the SPIO-HGNS-II and SPIO-HGNS-III samples exhibit an equilibrium temperature rise of 19.7 °C and 4.7 °C under the testing conditions, which is lower than that of SPIO-HGNS-I at the same Au concentration of 38 ppm (29.5 °C). Compared with the blank photothermal conversion test, where only a ~2.5 °C temperature increase is observed, the NIR triggered temperature rises on SPIO-HGNS-I and SPIO-HGNS-II are much higher, suggesting a strong photothermal conversion capacity. The temperature rise in the SPIO-HGNS-III sample was found to be small when compared to the other two samples. The reason for this reduced temperature rise in the SPIO-HGNS-III sample may be that the laser used for this sample is 945 nm. At this wavelength, the water absorption effect is much stronger than it at the wavelength around 800 nm. For example, the water absorption at 980 nm is 18 times higher than it at 808 nm [47]. Therefore, much more laser power can be absorbed by water during the temperature measurement of SPIO-HGNS-III, which would cause the observed small rise in temperature.

**Fig. 6.**
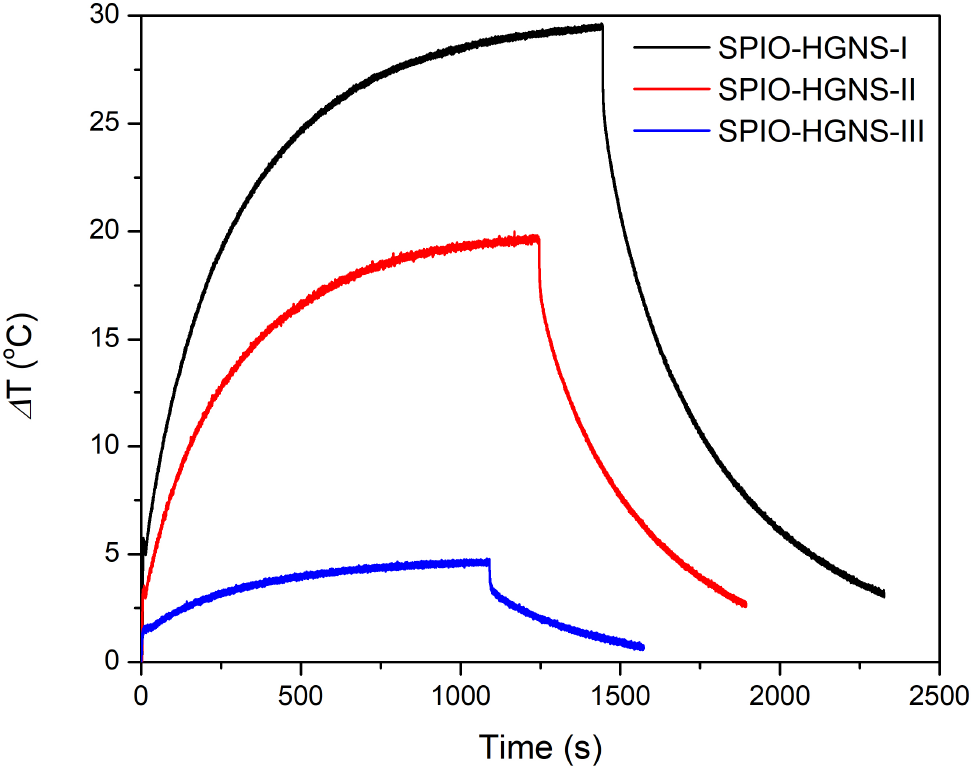
Temperature profile of SPIO-HGNS-I, II, and III at the Au concentration of 38 ppm.

### 3.4 The photothermal conversion efficiency evaluation

The photothermal conversion efficiencies of the as-synthesized SPIO-HGNS NPs at different sizes and concentrations were evaluated using our previously developed method that incorporates the experimentally measured temperature profiles into a FEM model [39]. In this FEM model, there are two volumetric heating sources (W/m^3^) considered in the system. The first heat source is from the heated thermocouple tip as a result of laser irradiation, defined as equation (1) [39]:

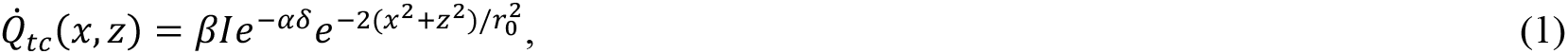

where *I* (W/cm^2^) is the laser intensity after passing the front cuvette wall and before entering the NPs solution; *δ* = 0.003 m is the distance between the thermocouple tip and the cuvette inner wall where the laser light penetrates; *α* is obtained from the experimentally measured absorbance value *A* using this equation: *α* = −ln(10^−*A*^)/*d*; *β* (m^−1^) is a value determined from the fitting of the simulated heating to the experimentally measured heating of DI water; *r_0_* is the beam radius = 0.001 m; the geometric center is defined at the thermocouple tip where *x* = 0, *z* = 0; *x* is defined along the vertical direction and *z* is defined along the horizontal direction, which is perpendicular to the laser direction.

The second heat source is from the laser irradiated NPs solution, defined as equation (2) [39]:

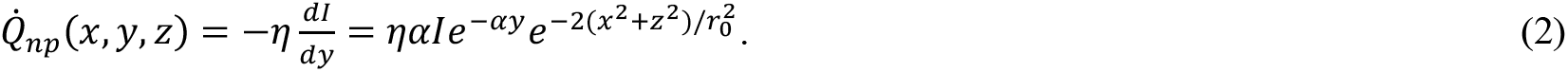

Where *η* is defined as the photothermal conversion efficiency, which is the fraction of the absorbed light that can be converted to heat; and *y* is defined along the direction of laser beam.

According to this FEM model, the photothermal conversion efficiency *η* of SPIO-HGNS NPs can be estimated using the following procedure:

First, the FEM simulation was performed to simulate the temperature profile of the pure DI water for the first 30 seconds of laser irradiation. For this case, the heating source was limited to the thermocouple tip. Thus, only equation (1) was used to define the heating source of the system. By comparing the simulated and experimentally measured temperature profile of water, the value *β* was determined to be 69681 m^−1^.

Next, the FEM simulation was performed to simulate the temperature profile of the SPIO-HGNS solution for the first 30 seconds of laser irradiation. For this situation, the heat sources include both the thermocouple tip and the NP solution. Thus, both equations (1) and (2) were used to define the heating sources of the system. An arbitrary *η* was assigned and the previously determined *β* was incorporated into the numerical modeling to simulate the temperature profile of the NP solution. The value of *η* was varied from 0 to 1 until a best fitting of the simulated to experimental temperature profile was obtained.

As shown in Fig. 7, during the very first 30 s, the simulated temperature profile is comparable to the experimental data. The value of the best-fitted photothermal conversion efficiency is summarized in Fig. 7. Photothermal conversion efficiencies of 0.55, 0.6, and 0.4 were obtained for SPIO-HGNS-I, II, and III, respectively. Notably, the photothermal conversion efficiency is slightly higher in SPIO-HGNS-II when compared with the other two samples. However, after 30 seconds, the experimentally measured temperature rise in SPIO-HGNS-I (5.9 °C) is higher than that of SPIO-HGNS-II (4.4 °C). The reason for this is that, even at the same concentration of 57 ppm, the light absorption value A of SPIO-HGNS-II at 855 nm is much higher than that of SPIO-HGNS-I at 820 nm. Therefore, the laser power that can reach the thermocouple tip position is much lower in the SPIO-HGNS-II sample than it is in the SPIO-HGNS-I sample. This lower laser power input at the thermocouple tip position causes the reduced temperature rise in SPIO-HGNS-II, despite the fact that the photothermal conversion efficiency of SPIO-HGNS-II is slightly higher than that of SPIO-HGNS-I. Similarly, there are also much fewer power inputs in the SPIO-HGNS-III sample, which further reduces the temperature rise to 2.7°C. Unlike the closed systems reported in the literature, such as a vacuum environment or a droplet liquid suspension [39], we use an open system to simplify the temperature measurement process, and there is energy transfer between the NP solution and surrounding environment. This energy loss may contribute to a smaller rise in temperature. Liquid evaporation may also affect the temperature rise and the determination of photothermal conversion efficiency. Additionally, the lower efficiency of SPIO-HGNS-III may come from the much stronger water absorption effect by the laser wavelength of 945 nm that was applied to this sample.

**Fig. 7.**
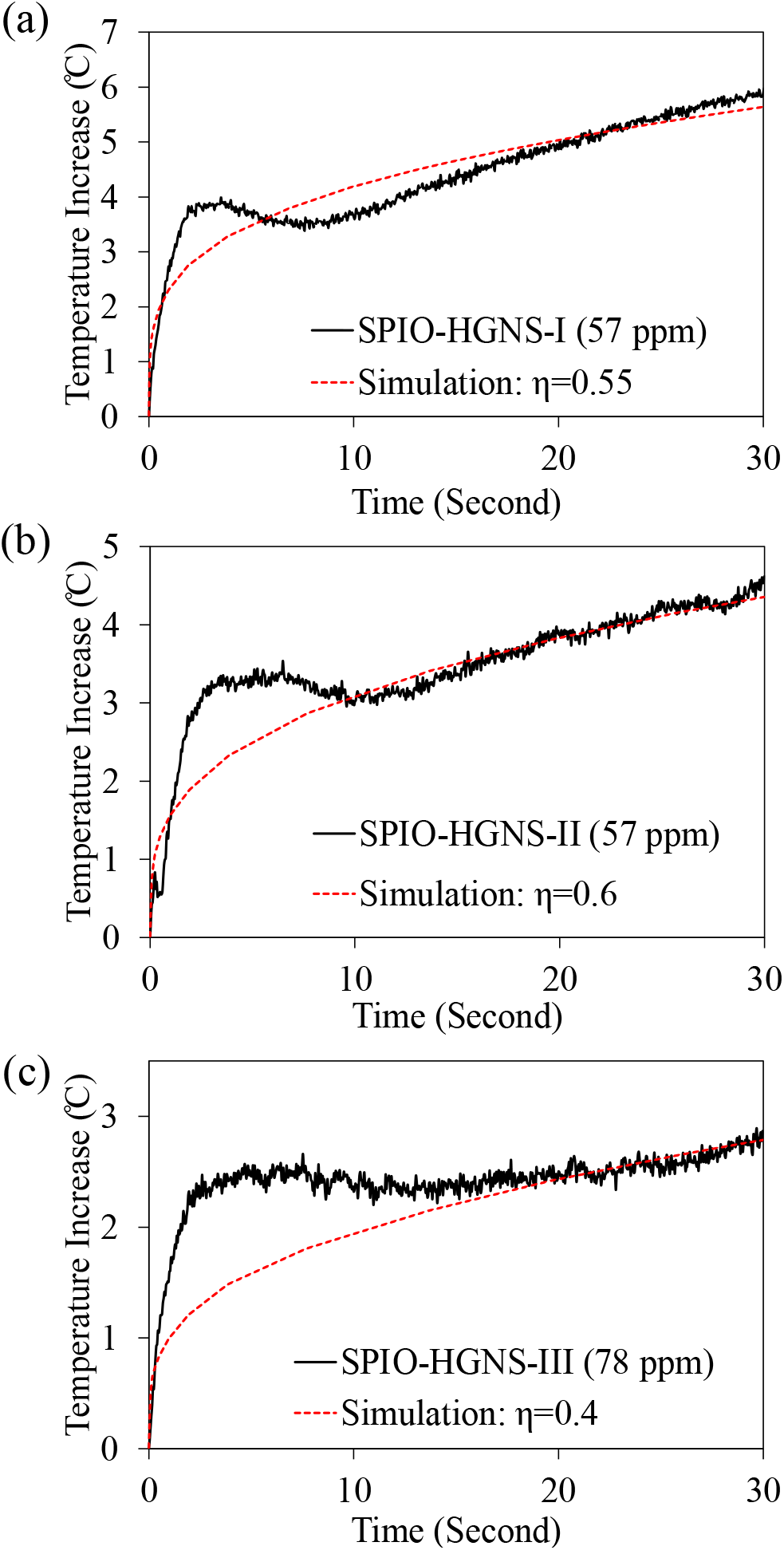
The measured vs. simulated temperature increases from COMSOL modeling for the very first 30 seconds for SPIO-HGNS-I, 57 ppm, 820 nm, light absorbance A=1.425, (b) SPIO-HGNS-II, 57 ppm, 855 nm, light absorbance A=1.868, (c) SPIO-HGNS-III, 945 nm, light absorbance A=2.481.

Compared to our SPIO-HGNS NPs that have a photothermal conversion efficiency of 0.6 (60%), literatures have previously reported Au nanoshells such as SiO_2_/Au and Au2S/Au have a photothermal conversion efficiency of 30%-39%, and 59%, respectively, while nanorods have a photothermal conversion efficiency of 55% [48, 49]. For pure HGNS, the photothermal conversion efficiency can go up to 90% [50]. However, these nanoshell structures possess limitations, such as absence of magnetic core and an inability to tune the SPR peak. This may restrict their biomedical applications, especially for brain diseases. Carbon nanocomposites are also reported to have outstanding photothermal conversion efficiencies as solar absorbers. However, as toxicity and biocompatibility are major concerns, the present knowledge around their safety is inadequate [51]. Attempts have been made to use Au nanorods for remote light-controlled drug delivery, such as Au nanorods with mesoporous silica/hydroxyapatite shell [52] and Au nanorods/microgel system [53]. However, the size of the hydrogel system is around 700 nm, which is too large for tumor uptake or applications in the brain.

Compared to the plasmonic nanostructures summarized in Table S3, our SPIO-HGNS possess benefits, such as tunable SPR peak at the NIR region and functional sizes within 10-100 nm, which enable enhanced tumor uptake, efficient blood brain barrier crossing and deeper brain tissue penetration [54, 55, 56, 57]. We have also shown that Au coating displays excellent biocompatibility and efficient cellular uptake [8, 32]. Additionally, another benefit is that the SPIO core enables magnetic targeting and therefore can enhance the accumulation of NPs at specific target sites and improve drug delivery performance [58, 59].

Under ultrafast laser irradiation, for all the SPIO-HGNS samples except the water case, a noticeable rapid rise, followed by a slight decrease in temperature, is observed. This phenomenon is not observed in continuous laser cases [39]. A possible reason for this is, when compared to continuous laser, the femtosecond pulsed laser leads to the nonlinear optical processes that trigger a rapid and sudden temperature rise in the plasmonic NPs. This localized and rapid temperature rise induces the formation of nanobubbles around plasmonic NPs [60], which may affect the characteristics of the temperature profile in NP solutions.

### 3.5 The plasmonic property analysis — The simulation of absorption cross-sections of SPIO-HGNS NPs

To further explore the plasmonic property of the as-synthesized SPIO-HGNS NPs, we employed a boundary element method proposed by Hohenester (MNPBEM toolbox) [61] to study the absorption properties of SPIO-HGNS NPs with fixed SPIO cores at 10 nm and different Au inner diameters and shell thicknesses. In our calculations, we assume that SPIO-HGNS NPs possess core-shell-shell structures with SPIO as the core, water as the interlayer, and HGNS as the outer layer. The refractive index for SPIO and water are 2.42 and 1.33, respectively. The refractive index for HGNS is simulated as a frequency-dependent function and is obtained from the MNPBEM toolbox. Fig. 8 (a) shows the absorption cross-sections of the multilayer SPIO-HGNS NPs for different HGNS inner diameters with fixed HGNS shell thickness of 6.7 nm. Fig. 8 (b) shows the absorption cross sections of the multilayer SPIO-HGNS NPs for different HGNS shell thicknesses with a fixed HGNS inner diameter of 40 nm. The results show that the increased inner diameter of HGNS leads to the longer wavelength (red-shifted) where the maximum absorption occurs, while the increased shell thickness of HGNS leads to the shorter wavelength (blue-shifted) where the maximum absorption occurs. The simulated results reveal how the change in HGNS inner diameter and shell thickness affect the position of maximum NIR light absorption, which provides strong guidance in designing plasmonic NPs with specific light absorption requirements. The results from TEM and light absorption spectra also demonstrate that the as-synthesized SPIO-HGNS NPs follow the same trend revealed from this simulation, in that the larger HGNS inner diameter induces the more red-shifted SPR wavelength of SPIO-HGNS NPs.

**Fig. 8.**
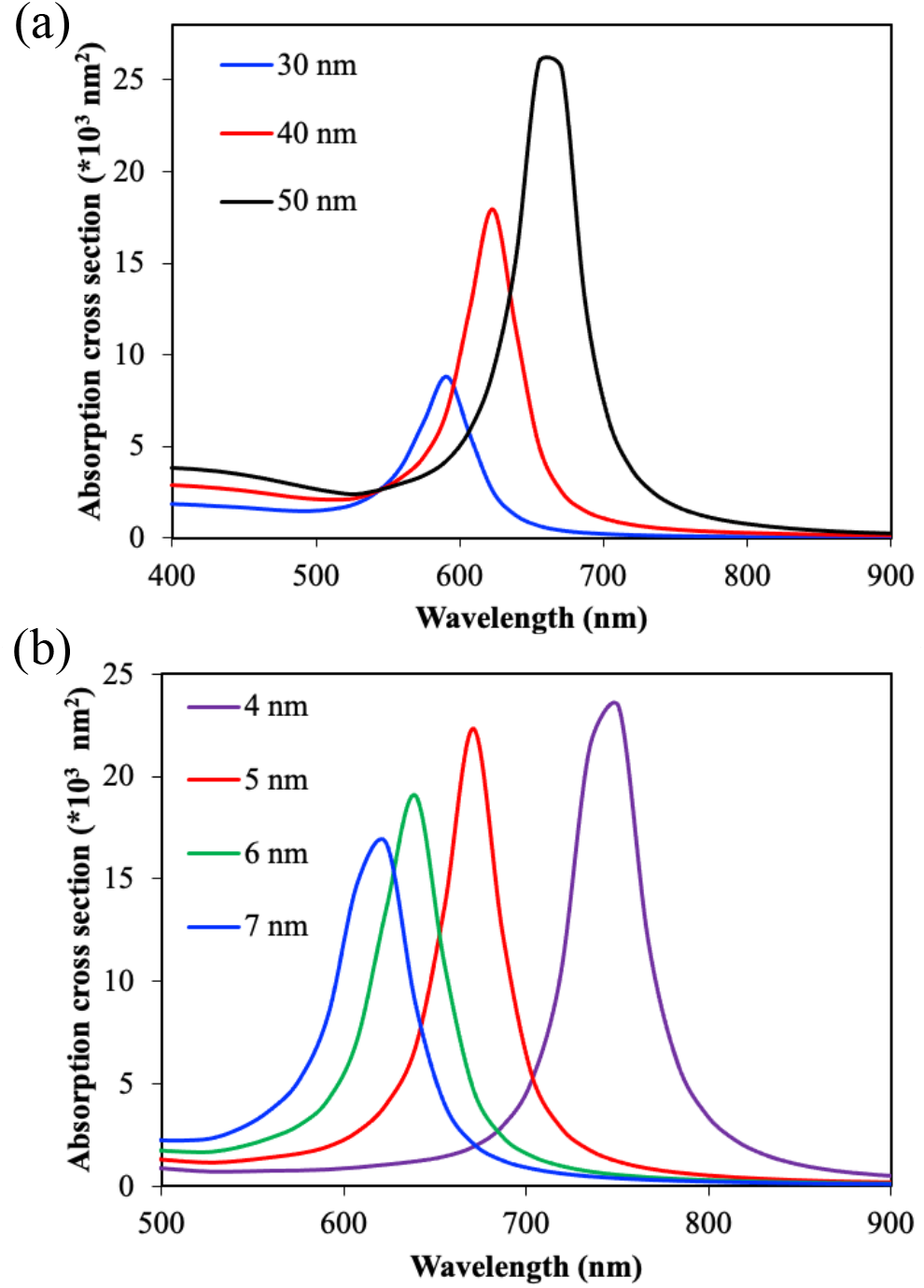
The absorption cross sections of SPIO-HGNS for (a) different HGNS shell inner diameters (30, 40 and 50 nm) with fixed HGNS shell thickness of 6.7 nm. (b) different HGNS shell thickness (4, 5, 6 and 7 nm) with fixed HGNS shell inner diameter of 40 nm.

## 4. Conclusions

In this work, we reported a novel method to synthesize SPIO-HGNS structures with tunable inner diameters and SPR peaks at the NIR region, which has a stronger tissue penetration capacity when compared to visible light. During the template-enabled galvanic replacement reaction, varying Ag^+^ concentrations were employed to control the inner diameter of HGNS. With the increasing Ag^+^ concentration, TEM imaging confirmed that the inner diameter of HGNS was enlarged. The existence of Au and Fe in the shell and core area was confirmed by Au111 lattice from TEM and EDS mapping of Au and Fe. The size quantification from TEM images showed that the synthesized SPIO-HGNS-I, SPIO-HGNS-II and SPIO-HGNS-III samples have similar shell thicknesses of 6.7, 6.5 and 7.0 nm and increased inner diameters of 38.7, 39.4 and 40.7 nm. ICP results estimated the mole ratio of Au/Fe to be 1.1, 2.2, and 5.9 for SPIO- HGNS-I, SPIO-HGNS-II and SPIO-HGNS-III, respectively. The light absorbance spectra revealed that, with the increasing inner diameter, SPIO-HGNS-I, SPIO-HGNS-II and SPIO-HGNS-III had red-shifted SPR peaks at 820 nm, 855 and 945 nm, respectively. The absorption cross-section calculation confirmed this same trend using models of SPIO-HGNS with different inner diameters and shell thicknesses. The as-synthesized SPIO-HGNS samples were highly efficient photothermal converters with maximum temperature rises of 29.5 °C, 19.7°C and 4.7 °C after 1000 s of laser irradiation and photothermal conversion efficiencies of 0.55, 0.6 and 0.4 for SPIO-HGNS-I, SPIO-HGNS-II and SPIO-HGNS-III samples, respectively. This suggests their strong potential for photothermal therapy. In the future, we will develop such nanomedicine for deep brain stimulation using low power NIR light.

## Availability of data and material

The datasets generated during and/or analyzed during the current study are available from the corresponding author on reasonable request.

## Authors’ contributions

The manuscript was written through contributions of all authors. All authors have given approval to the final version of the manuscript. Muzhaozi Yuan and Xuhui Feng contributed equally.

## Statements and Declarations

The authors declare that they have no conflict of interest.

## Acknowledgments

This work was kindly supported by the United States National Science Foundation (Award # CMMI 1851635, Y.W.; Award # ECCS 2021081, Y.W. and Y.L.). The use of Material Characterization Facility at Texas A&M University was acknowledged. The authors would like to thank MacKenzie Harnett for her assistance on manuscript revision and proofreading.

