## Supplemental Information for "Superparamagnetic iron oxide enclosed hollow gold nanostructure with tunable surface plasmon resonances to promote near-infrared photothermal conversion"

For

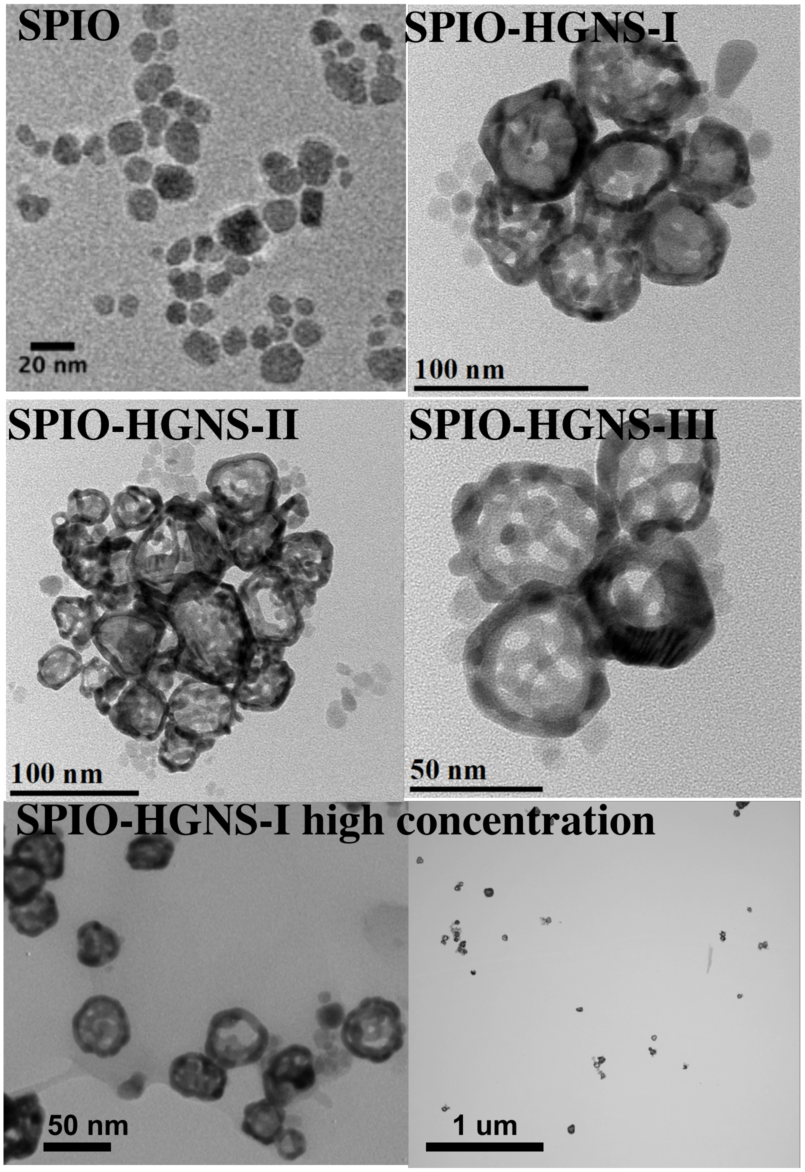


Figure S1. TEM images of SPIO NPs, SPIO-HGNS-I, SPIO-HGNS-II, SPIO-HGNS-III, and SPIO-HGNS-I NPs at high concentration.

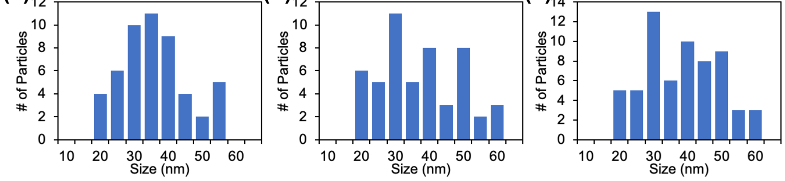

Figure S2. The size distribution of (a) SPIO-HGNS-I, (b) SPIO-HGNS-II, (c) SPIO-HGNS-III based on more than 50 NPs.

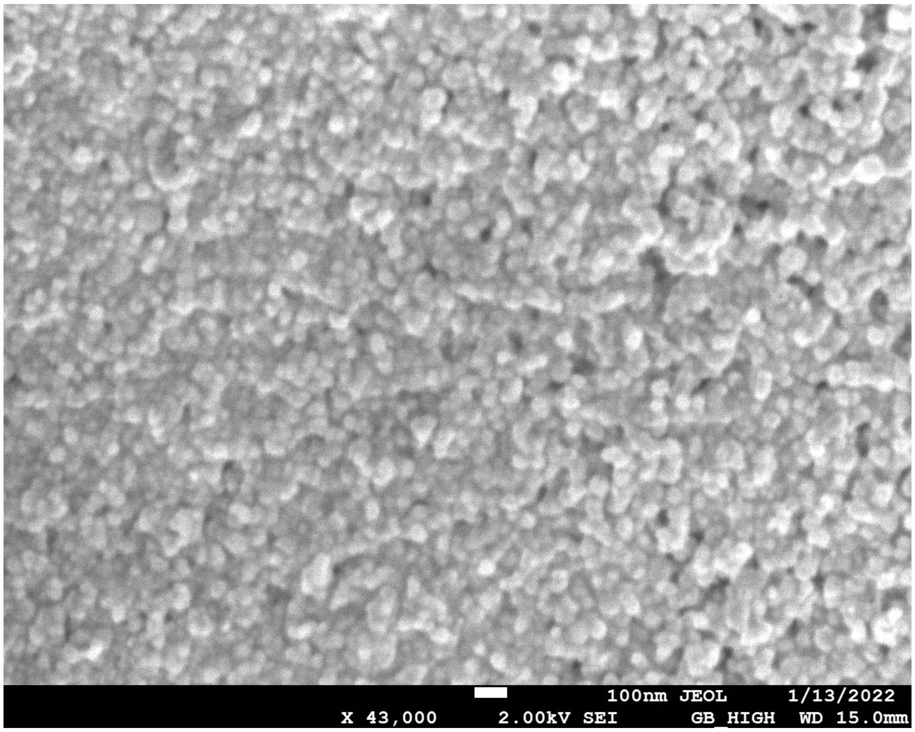


Figure S3. High resolution SEM image of SPIO-HGNS-I NPs.

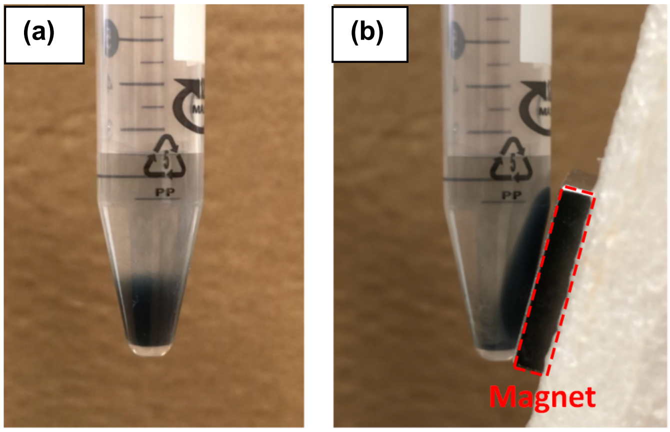


Figure S4. Photographs of the magnetic separation of SPIO-HGNS NPs (a) before and (b) after the application of magnet for 5 seconds, showing the accumulation and movement of SPIO-HGNS NPs towards the direction of magnetic field.

Table S1. ICP results showing the element concentration and the mole ratio of Au/Fe for SPIO-HGNS-I, II, and III NPs.

| Sample name | Fe (ppb) | Au (ppb) | Au/Fe in mole ratio |
|---|---|---|---|
| SPIO-HGNS-I | 17.3 ± 6.5 | 66.5 ± 1.7 | 1.1 |

| | | | |
|---|---|---|---|
| SPIO-HGNS-II | 9.5 ± 1.2 | 73.1 ± 4.0 | 2.2 |
| SPIO-HGNS-III | 3.0 ± 1.1 | 62.3 ± 1.2 | 5.9 |

Table S2. Zeta potential and hydrodynamic diameter of SPIO-HGNS NPs functionalized with thiol-PEG

| | SPIO-HGNS | Thiol-PEG modified SPIO-HGNS |
|---|---|---|
| Zeta potential (mV) | -44.5 ± 1.4 | 30.3 ± 0.3 |
| Hydrodynamic diameter (nm) | 211.8 ± 14.3 | 160.8 ± 2.4 |

Table S3. Summary of literatures reported plasmonic structures

| Materials | Size (nm) | Photothermal efficiency | SPR peak position (nm) | Benefits | Limitations |
|---|---|---|---|---|---|
| $SiO_2$ core/Au [1, 2] | 77 | 30-39% | 796-815 | SPR peak at NIR region | Lack of magnetic targeting and SPR tuning |
| $Au_2S$ core/Au [2] | 25 | 59% | 800 | SPR peak at NIR region | Lack of magnetic targeting and SPR tuning |
| HGNS [3] | 70-90 | 90% | 790 | High photothermal conversion, SPR peak at NIR region | Lack of magnetic targeting and SPR tuning |
| Au nanorod [2] | 44 in length, 13 in diameter | 55% | 780 | Tunable two distinct SPR peak at visible and NIR region | Lack of magnetic targeting, synthesized along with toxic agents which reduce the biocompatibility |
| Carbon composite[4] | 1-3 um | 89% | Solar absorption | High photothermal conversion | Lack of magnetic targeting, too large size, inadequate toxicity data. |

**The exploration of the relationship between SPR peak and size of HGNS**

To further explore the relation between SPR peak and size of HGNS, we summarized the results from previous paper and simulated their SPR peak, as shown in Table S4. It shows that for some of the HGNS structures, the measured SPR peak is very close to the simulated SPR peak, while for other HGNS structures, the measured SPR peak is greatly redshifted compared with the simulated SPR peak. The results suggest that the SPR peak of HGNS may be highly dependent to the surface morphology . Some references mentioned that the red-shifted SPR peak of HGNS with rough surface is induced by the strong interacted local electric fields from the neighboring bumps at the surface of HGNS shell, similar to the properties of plasmonic NPs cluster [5].

Table S4. Summary of HGNS size and SPR peak

| Type of NPs | Synthesis method | Inner diameter (nm) | Shell thickness (nm) | Measured SPR peak (nm) | Simulated SPR peak by our modeling (nm) |
|---|---|---|---|---|---|
| HGNS [6] | galvanic exchange on $Co_2B$ templates | 35.2 | 6.4 | 635 | 607 |
| | | 38.2 | 4.9 | 700 | 655 |
| | | 39.6 | 4.2 | 765 | 719 |
| HGNS [3] | galvanic exchange on $Co_xB_y$ templates | 46.0 | 5.0 | 660 | 718 |
| | | 46.0 | 12.0 | 720 | 591 |
| | | 46.0 | 22.0 | 760 | 559 |
| HGNS [7] | galvanic exchange on silver templates | 26.2 | 3.4 | 820 | 655 |
